# HD mutation results in a dominant negative effect on HTT function

**DOI:** 10.1101/2023.06.26.543767

**Authors:** Tiago L. Laundos, Shu Li, Eric Cheang, Riccardo De Santis, Francesco M. Piccolo, Ali H. Brivanlou

## Abstract

Huntington’s disease (HD) remains an incurable and fatal neurodegenerative disease long after CAG-expansion mutation in the huntingtin gene (HTT) was identified as the cause. The underlying pathological mechanism, whether HTT loss of function or gain of toxicity results from mutation, remains a matter of debate. In this study, we genetically modulated wild-type or mutant HTT expression levels in isogenic human embryonic stem cells to systematically investigate their contribution to HD-specific phenotypes. Using highly reproducible and quantifiable *in vitro* micropattern-based assays, we observed comparable phenotypes with HD mutation and HTT depletion. However, halving endogenous wild-type HTT levels did not strongly recapitulate the HD phenotypes, arguing against a classical loss of function mechanism. Remarkably, expression of CAG-expanded HTT in non-HD cells induced HD-like phenotypes akin to HTT depletion. By corollary, these results indicate a dominant negative effect of mutated HTT on its wild-type counterpart. Complementation with additional copies of wild-type HTT ameliorated the HD-associated phenotypes, strongly supporting a classical dominant negative mechanism. Understanding the molecular basis of this dominant negative effect will guide the development of efficient clinical strategies to counteract the deleterious impact of mutant HTT on the wild-type protein.

## Introduction

Huntington’s disease (HD) is a heritable, fully penetrant, autosomal neurodegenerative disease caused by a CAG trinucleotide expansion at the 5’ end of the Huntingtin (HTT) gene (Macdonald 1993), leading to an expanded poly-glutamine (poly-Q) stretch at the N-terminal portion of the encoded HTT protein. Expansions above 35CAG are in the pathological range (Kay et al. 2016), and age of clinical onset correlates with CAG repeat length, with larger expansion leading to earlier onset of motor symptoms (Andrew et al. 1993; M. Duyao et al. 1993). Due to selective neuronal vulnerability, HD mutation results in extensive loss of striatal medium spiny neurons and cortical neurons, a disease hallmark evident in postmortem HD patient brains (Aylward et al. 2011; Halliday et al. 1998; Ross and Tabrizi 2011). Three decades since the genetic cause was identified, the molecular mechanism by which a monoallelic mutation results in disease onset in adult is yet to be fully understood, and a cure or treatment to stop the progression of the disease is currently nonexistent.

HTT protein is ubiquitously expressed in all tissues from the early embryo onwards and despite its function being still largely elusive, it has been suggested to participate in a variety cellular process, such as vesicle transport (Caviston et al. 2007), ciliogenesis (Haremaki, Deglincerti, and Brivanlou 2015), cell survival (Zhang et al. 2006), and establishment of polarity (Galgoczi et al. 2021). Animal models showed that lack of HTT expression is embryonically lethal at day E7.5 (M. P. Duyao et al. 1995) and no reports of healthy human patients devoid of HTT have been presented. Furthermore, mouse forebrain-specific knockout of HTT results in progressive neurodegeneration (Dragatsis, Levine, and Zeitlin 2000), suggesting that HTT is required both for proper embryonic development and for maintenance of the neuronal population.

While several mouse models have been generated to recapitulate some of the hallmarks of HD pathophysiology, such as loss of striatal neurons and motor dysfunction (reviewed in (Figiel et al. 2012)), they pose limitations due to species differences and lack of complete phenotype. Alternatively, *in vitro*, it has been shown that HD mutation results in neuronal dysfunction, such as altered axonal transport of BDNF, abnormal mitochondrial dynamics, and mitotic spindle orientation (Gauthier et al. 2004; C. Lopes et al. 2016; Shirendeb et al. 2012). However, the severity of these phenotypes associated with HD mutation are challenging to quantitatively attribute(relate) to the various CAG lengths and HTT expression levels, hindering clarification of the mechanism behind HD clinical presentation. Thus, the consequences of HD mutation on the physiological function of HTT remain mostly unknown.

Our laboratory has previously generated a series of micropattern-based *in vitro* assays, which unveiled novel and subtle HD-signature phenotypes in standardized and reproducible spatially-confined cultures with defined geometry that are well-suited for direct quantitative measurements of the spatial organization of cells. Besides revealing new HD-associated phenotypes, these assays proved to be highly sensitive and specific by displaying increased phenotypic severity correlating with CAG domain expansion level (Ruzo et al. 2018; Galgoczi et al. 2021; Haremaki, Deglincerti, and Brivanlou 2015; Piccolo et al. 2022). Additionally, the quantification of these phenotypes proved that lack of HTT expression invariably recapitulates the signatures observed in CAG expanded lines with the highest degree of severity, suggesting that CAG expansion mutation might result in the progressive loss of HTT functions.

In this study we aim to systematically assess the effect of HD mutation on HTT function and directly test the hypothesis that the phenotypes observed are due to loss of HTT function, employing two alternative micropattern-based assays: 1) a fast signaling readout assay, consisting of 1 hour of Activin A stimulation of confluent pluripotent cells so that in control condition only cells at the edge of the colony are activated, while the HD-signature consists in the activation of most cells regardless of their geometrical positioning (Galgoczi et al. 2021); 2) a long-term assay, neuruloid formation in 7 days culture, which models the self-organization of major ectodermal lineages during human neurulation and where the HD-signature phenotype results in failed compaction of the central neural ectodermal domain and in the reduction of the Neural Crest linage (Haremaki, Deglincerti, and Brivanlou 2015). These are highly reproducible standardized culture systems that allow accurate quantitative assessment of small phenotype changes.

Using these two assays, we show that halved level of HTT expression results in a very mild HD-like phenotype, significantly less severe than in HD hESCs. Moreover, we show that the expression of a mtHTT transgene in WT cells containing a full dose of endogenous wtHTT results in the induction of a HD phenotype, mimicking the phenotypes observed in HD monoallelic mutation and in HTT^-/-^ cells. This indicates that mtHTT has a deleterious effect on wtHTT activity. Moreover, genetic complementation of HD lines with exogenously expressed full length wtHTT can rescue their phenotypes in micropattern-based assays, but not fully, suggesting HD mutation results in a Dominant Negative effect.

## Results

### Lowering HTT expression induces HD-signature phenotypes

To directly assess the consequences of CAG expansion mutation on HTT physiological function we compared the phenotypes of a HD hESC line, where one HTT allele carries a 72CAG expansion mutation (HTT^WT/MT^, HD), with isogenic lines expressing half of the physiological level of HTT (HTT^WT/-^, heterozygote) or none (HTT^-/-^, hereafter referred as HTT-KO) that we previously generated ((Ruzo et al. 2018)) (Figure 1A, B and C). In wild-type (HTT^WT/WT^, WT) hESC confluent micropattern culture, following 1 hour stimulation with activin A (Figure 1D), cells at the edge of the colonies were activated and accumulate SMAD2/3 in the nucleus in response to apical activin A induction (Figure 1E, F), while cells at the center of the micropattern colonies did not show nuclear translocation of SMAD2/3. On the other hand, and as previously shown (Galgoczi et al. 2021), micropattern colonies formed by HD lines lost the spatial restriction of sensitivity to activin A, allowing the response to the morphogen in cells residing at the center region of the micropatterns (Figure 1E, F). In this experimental setting, the heterozygous line (HTT^WT/-^) showed a very subtle increase in SMAD2/3 nuclear signal in the center of the colony and overall significant increase in the total number of SMAD2/3+ nuclei (Figure F, G), while lack of HTT expression (HTT-KO) resulted in the nuclear translocation of SMAD2/3 throughout the micropattern colony. This indicates that reduced level of HTT expression results in an HD-signature phenotype, albeit significantly milder than complete lack of HTT expression or monoallelic HD mutation.

**Figure 1.**
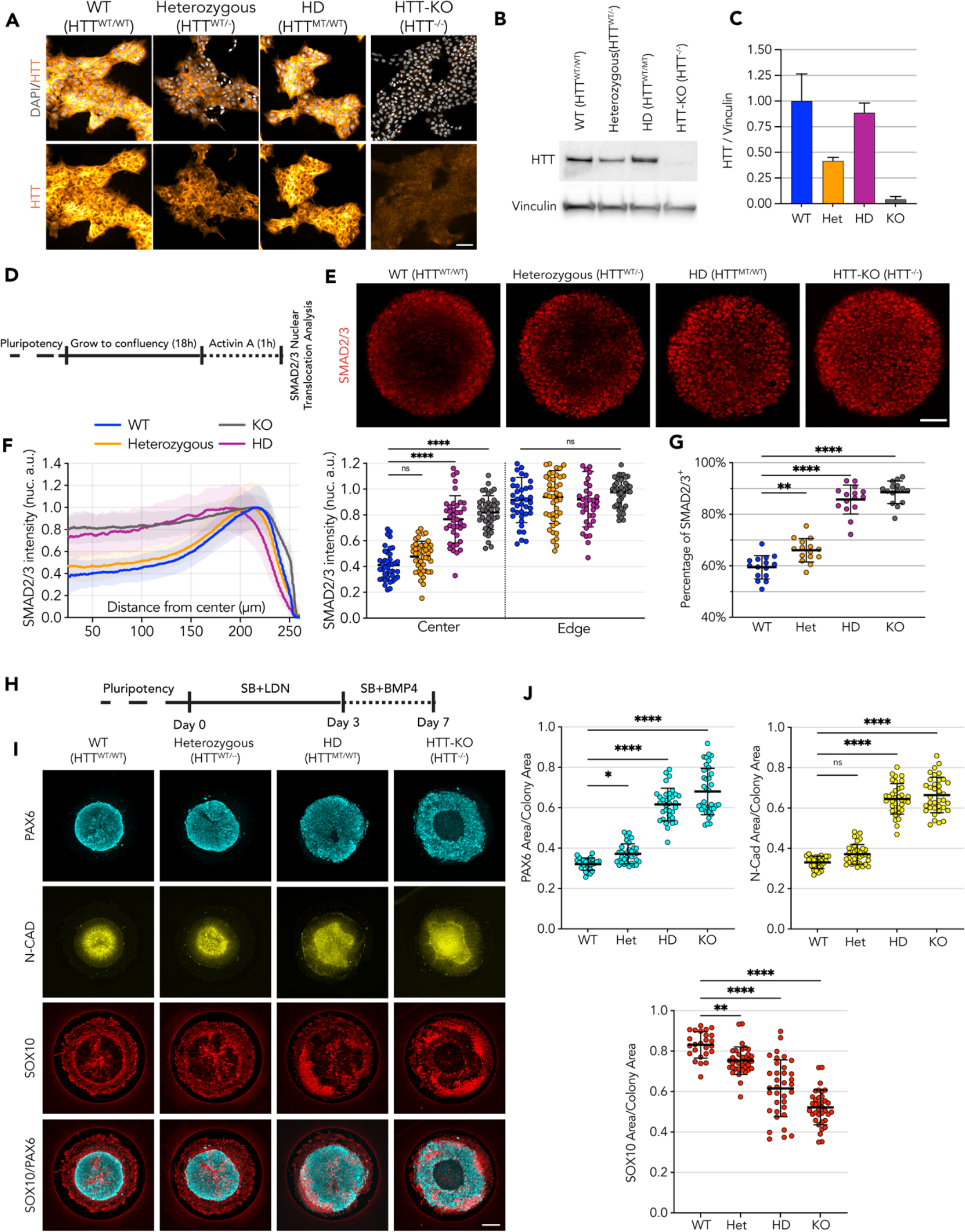
HD-signature phenotype in micropattern cultures show loss of HTT function. **A** Immunofluorescence detection of HTT of pluripotent embryonic stem cells illustrates reduction of HTT protein in heterozygous cell line (D7F7 antibody). HD cell line genetic makeup is HTT^72CAG/20CAG^. **B** Immunoblot against HTT shows reduced HTT protein expression in heterozygous cell line (MAB2166 antibody). Vinculin shown as loading control. **C** Immunoblot signal quantification confirms halving of HTT amount compared to WT and HD lines (n=3 independent experiments). **D** Schematics of Activin A assay procedure performed on 500µm circular micropatterns. **E** Immunofluorescence imaging shows SMAD2/3 nuclear translocation upon Activin A stimulation, limited to the edges on WT and Het colonies with wider to full induction on the colony center HD and HTT-KO colonies, respectively. **F** Mean radial intensity profile of nuclear SMAD2/3 shows insensitivity to Activin A at the colony center in Het cell lines is partially disrupted by lower HTT levels, with patches of activated cells in the colonies center, in contrast with HD cell line which presents a phenotype comparable to HTT-KO with SMAD2/3 translocation induced throughout the micropatterned colonies (scatter plot displays the mean SMAD2/3 nuclear intensity for each colony at center:25-10µm and edge:175µm-225µm; WT n=41, Het n=44, HD n=39, KO n=41) **G** Quantification of SMAD2/3^+^ nuclei numbers reveals an increase of the fraction of SMAD2/3^+^ in Het cell lines, in line with a mild HD-like phenotype (n=15 colonies per condition). **H** Schematics of the Neuruloid induction assay, performed on 500µm circular micropatterns. **I,J** HTT halving in Het cell line results in enlarged neuroectodermal domain (PAX6 and NCAD area) and a reduction of the neural crest lineage (SOX10 area). HD shows a significantly worsened phenotype than Heterozygous cell line suggesting monoallelic CAG expansion on HD cell line does not result in from a classical loss of function of HTT upon mutation (WT n=25, Het n=33, HD n=34, KO n=36). Groups were compared using one-way ANOVA followed by Dunnett’s *post-hoc* test for correction of multiple comparisons to the WT control (* p<0.05, ** p<0.01, *** p<0.001, ****p<0.0001). All values are presented as mean ± SD. Scale bar: 100µm.

The consequences of lower HTT expression were also explored in the neuruloid formation assay (Figure 1H, Haremaki et al). Here, HD hESC (HTT^WT/MT^) showed a loosened central neuroectodermal domain, measured as an increased PAX6^+^ area and N-Cad lumen, along with a contraction of the Neural Crest lineage, quantified as a decrease of the area occupied by SOX10^+^ cells (Figure 1 I, J) (Haremaki et al). The neuruloids derived from the heterozygous (HTT^WT/-^) line, that express half of the physiological HTT level, displayed an HD-like phenotype in all three measurements. However, the extent of this phenotype was less severe than the phenotype observed in HD lines, which grossly failed to compact the central neuroectodermal area, a phenotype only worsened by complete lack of expression of HTT protein, as in HTT-KO cell line (Figure 1 F and G).

Taken together, these results show that reduced expression of HTT protein recapitulates aspects of the phenotypes observed in HD samples, suggesting that HD mutation might result in a loss of HTT function. However, when compared to the more severe HD phenotype, the milder HD-like phenotypes displayed by heterozygote cells provides evidence supporting the idea that monoallelic CAG expanded HTT might also exert deleterious activity that results in more severe phenotypes.

### CAG-Expansion affects HTT function

To dissect the effect of HD mutation on HTT function and its role in these platforms, we designed a transgenic system that allows us to modulate the expression of wild-type and mtHTT proteins in the various isogenic backgrounds of the RUES2 HD allelic series (Ruzo et al). For this purpose, we engineered two Piggybac vectors containing the coding sequence of full length human HTT carrying a normal trinucleotide stretch (wtHTT, 20CAG) or an expanded CAG stretch (mtHTT, 56CAG). At the 3’ end of the HTT synthetic gene, an in-frame sequence coding a Glycine-Serine flexible linker and the HaloTag coding sequence were introduced, resulting in a C-terminal tagging of the transgenic full-length HTT proteins, which are expressed under a constitutive promoter (Figure 2A). These vectors (wtHTT or mtHTT) were stably transfected into the HTT-KO cell line, and successful in-frame full length expression was verified by immunoblot, with detection of the HTT as well as the HaloTag amino acid sequences, confirming that the levels of expression of the two transgenes in the cell lines generated were equivalent (Figure 2B, Sup Figure 3).

**Figure 2.**
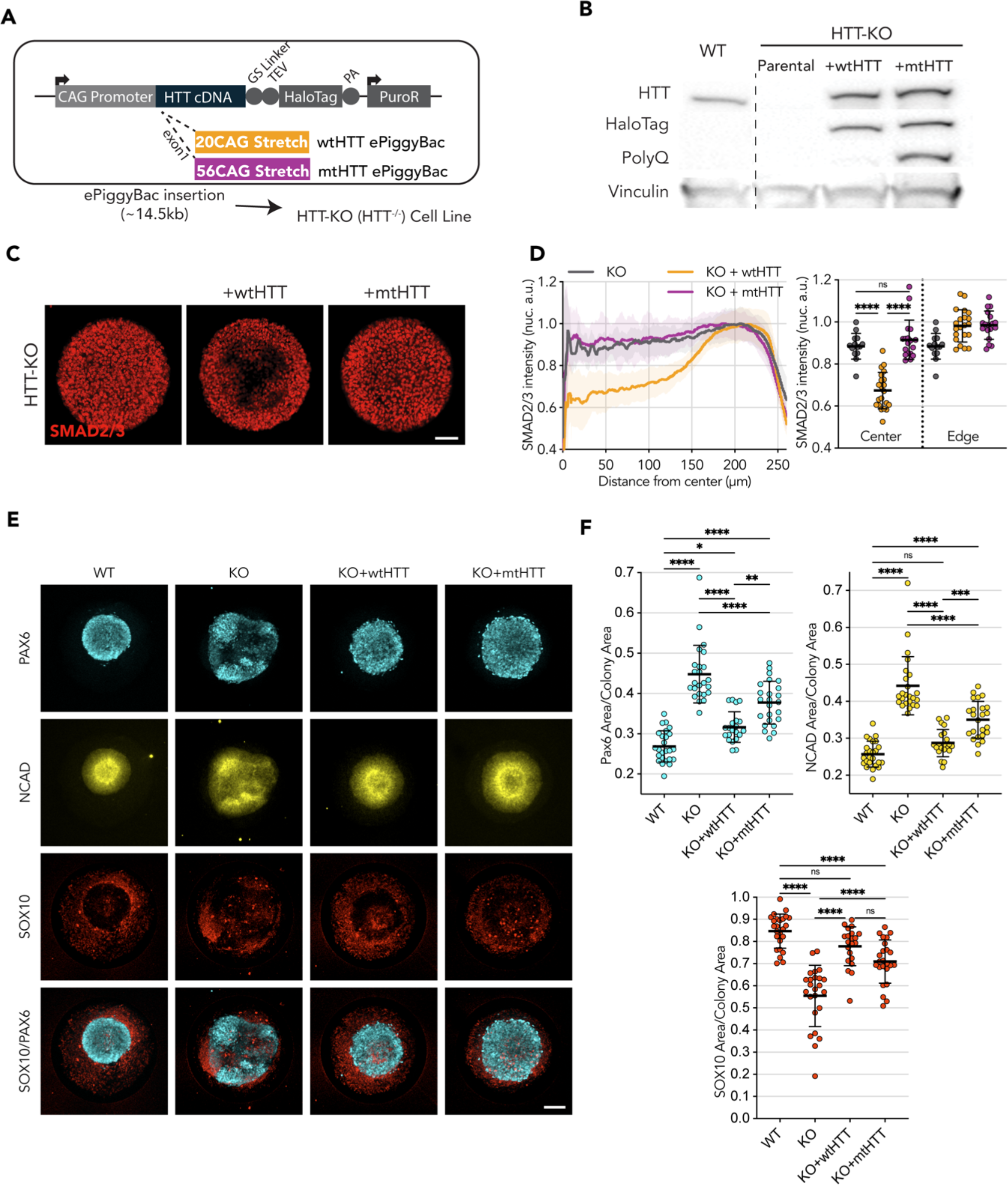
Wild type HTT, but not mutant, restores endogenous HTT function. **A** Puromycin selectable ePiggyBac transposon carrying the full length HTT coding DNA sequence, to be expressed under the CAG promotor, and fused at the C-terminal with HaloTag spaced by a Glycine-Serine Linker for added flexibility. Two vectors were built to contain a 20CAG (wt) or 56CAG (mt) stretch on exon 1. The ∼14.5kb transposon was inserted in an HTT-KO cell line with ablated endogenous HTT expression. **B** Immunoblot against HTT (MAB2166) shows exogenous HTT protein expression in KO+wtHTT and KO+mtHTT; HTT exogenous expression lines also present HaloTag epitope around HTT size (∼350kDa) showing tag remains fused; mutant HTT transposon in KO+mtHTT is detected with anti-PolyQ specific antibody (MW1). Vinculin shown as loading control. **C** Expression of exogenous wtHTT restores edge restriction in response to activin A in HTT-KO, with absence of SMAD2/3 nuclear translocation at the colony center (KO+wtHTT), an effect that is not observed with the mutant counterpart. **C,D** Mean radial intensity profile of nuclear SMAD2/3 confirms this effect (HTT-KO n=14, KO+wtHTT n=21, KO+mtHTT n=18). **E, F** Exogenous expression of wtHTT, and to a lesser extent of mtHTT, result in a reduction of the neuroepithelial domain (PAX6 and NCAD) as well as an increase in neural crest domain (SOX10). (WT n=25, HTT-KO n=24, KO+wtHTT n=21, KO+mtHTT n=24). Groups were compared using one-way ANOVA followed by Tukey’s *post-hoc* test for correction of multiple comparisons (* p<0.05, ** p<0.01, *** p<0.001, ****p<0.0001). All values are presented as mean ± SD. Scale bar: 100µm.

**Figure 3.**
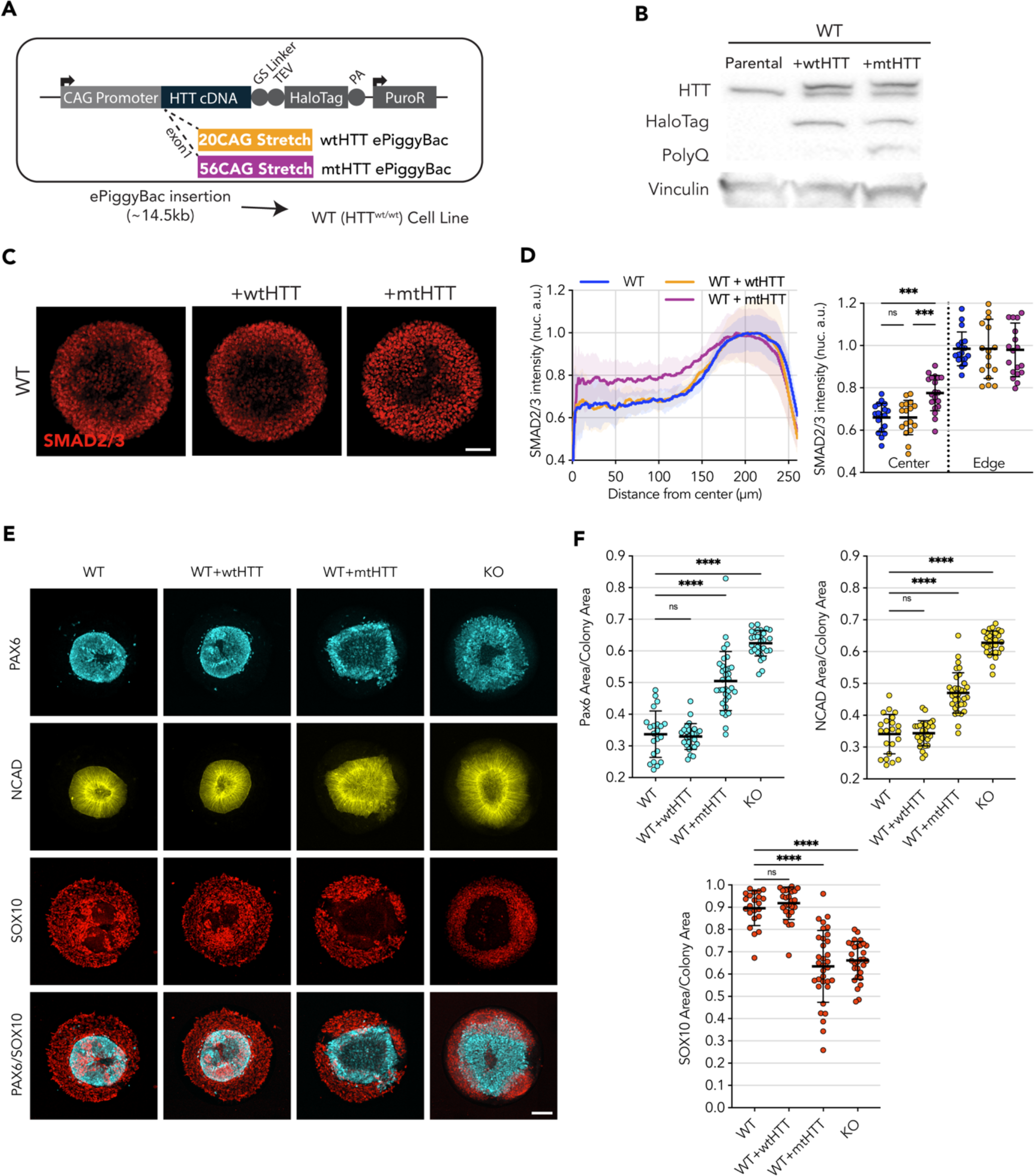
Transgenic expression of full length mt-HTT induces HD phenotype in WT cell lines. **A** Puromycin selectable ePiggyBac transposon carrying the full length HTT coding DNA sequence and containing either 20CAG (wt) or 56CAG (mt) stretch on exon 1 was inserted in an WT cell line, which originally expresses wtHTT protein at endogenous levels. **B** Immunoblot against HTT (MAB2166) shows endogenous expression of HTT across the 3 lines (∼350kDa); WT+wtHTT presents a second band corresponding to the inserted transgene, at higher molecular weight than wtHTT due to the fusion to HaloTag(+36kDa); WT+mtHTT also displays an additional transgene band, though at higher molecular weight than the former wtHTT transgene due to the addition of the expanded poly-glutamine stretch on the mutant version (additional 36 Glutamines; +12kDa). HaloTag expression is confirmed for both exogenous HTT expression cell lines as well as specificity of PolyQ expansion on WT+mtHTT cell line (MW1 antibody). Vinculin shown as loading control. **C, D** Mean radial intensity profile of nuclear SMAD2/3 reveals expression of exogenous mtHTT in WT cells disrupts edge sensing restriction to activin A stimulation (1hr), resulting in SMAD2/3 nuclear translocation in central areas of the micropattern in an HD like phenotype, while wtHTT overexpression did not impact the WT radial profile (WT n=18, WT+wtHTT n=17, WT+mtHTT n=19). **E** Neuruloid formation assay shows insertion of mtHTT leads to increased PAX6/NCAD neuroectodermal domain In a HD like Phenotype. **F** Quantification of confirms the increased PAX6 and NCAD domain in WT cells presenting the mtHTT transgene, as well as a reduction in the SOX10^+^neural crest domain to a level seen in HTT-KO neuruloid suggesting a Dominant negative effect of mtHTT on WT cells. Groups were compared using one-way ANOVA followed by Dunnett’s *post-hoc* test for correction of multiple comparisons to the WT control (* p<0.05, ** p<0.01, *** p<0.001, ****p<0.0001). All values are presented as mean ± SD.

To determine whether HaloTag C-terminal fusion affects HTT activity, we tested for its ability to rescue the phenotype of HTT-KO hESCs in the two micropatterned-based assays. First, we performed the 1hr activin A induction protocol where expression of wtHTT-HaloTag was able to rescue the phenotype of HTT-KO cells, resulting in a SMAD2/3 nuclear translocation restricted to edge of the micropatterns (Figure 2C, D). In contrast, expression of mtHTT-HaloTag failed to rescue the HD-like phenotype of edge restricted SMAD2/3 activation (Figure 2C, D). Similarly, exogenous expression of wtHTT-HaloTag was also able to rescue the HD-like phenotype of HTT-KO hESC in the neuruloid assay, allowing for proper compaction of the central neuroectodermal region, measured as reduced area of the PAX6^+^ domains and the NCAD^+^ lumina (Figure 2E and F), as well as rescue of the Neural Crest SOX10^+^ area. However, exogenous expression of full-length mtHTT in HTT-KO hESC only led to partial rescue of its HD-like phenotype shown by a less pronounced reduction in the area PAX6^+^ and NCAD+ domains, and milder increase in SOX10^+^ domain, indicating that in this more complex model, mutant HTT retains some activity for the at least a partial rescue of the phenotype in cells devoid of HTT expression.

These results indicate that CAG-expansion might directly impair the function of HTT protein.

### mtHTT expression is sufficient to induce HD-like phenotype in Wild-Type hESCs

After assessing the functionality of wtHTT and mtHTT transgenic proteins in HTT-KO hESCs, we exogenously expressed them in WT hESCs to directly test the effect of mutant HTT on the two micropattern assays (Figure 3A and B). As a control, overexpression of wtHTT in WT hESCs did not affect the ability of WT cells to spatially restrict sensitivity to Activin A upon 1hr of stimulation, with nuclear translocation of SMAD2/3 limited to the edges of their colonies (Figure 3C, D). Similarly, WT hESCs overexpressing wtHTT was able to generate neuruloids that properly form a compact central neuroectodermal structure (PAX6/NCAD), which was surrounded by a well-formed Neuroectodermal structure (SOX10) (Figure 3E and F).

On the other hand, exogenous expression of mtHTT profoundly affected the ability of WT hESC cells to perform in the two micropatterns assays. Upon 1 hour of Activin A stimulation, large patches of cells exhibiting nuclear translocation of SMAD2/3 appeared toward the center of the colony, reminiscent of the HD phenotype previously shown (Figure 3C and D). Likewise, in the neuruloid assay, a similar effect was observed with the micropatterns presenting grossly misshapen and enlarged central domains (as shown by PAX6 and N-CAD signal) as well as a reduction of the neural crest lineage (SOX10), recapitulating the phenotype previously characterized for HD line (Figure 3E and F).

These observations show that the expression of a mutant form of HTT in a fully competent WT (HTT^WT/WT^) background is sufficient to elicit HD-like phenotypes. This indicates that CAG expansion mutation on the HTT gene results in some kind of detrimental activity that might interfere with physiological processes performed by endogenous HTT in WT cells.

### Genetic complementation with full length WT HTT partially rescues the phenotypes of HD lines

Since exogenous expression of mtHTT induces HD-like phenotypes in WT hESCs, while only partially rescuing the HD-like phenotypes of HTT-KO hESCs, we hypothesized that CAG expansion mutation might result in a reduced function of HTT protein and in some gained activity that somehow interferes with the physiological function of wild-type proteins. To test this hypothesis, we sought to genetically complement an HD hESC line, carrying endogenous monoallelic 72CAG expansion mutation (HTT^WT/MT^), with our construct encoding full length wtHTT-HaloTag protein and determine whether excess of wtHTT protein would rescue its HD-signature phenotypes (Supplementary Figure S4A). Stable expression of HTT-HaloTag was verified by immunoblot and HaloTag detection. (Supplementary Figure S4B, C).

Differently from the HTT-KO cell line, complementation of HD hESC lines with wtHTT-HaloTag partially, but significantly rescued the HD phenotype on the Activin A assay (Supplementary Figure S4D,E). Likewise, on the neuruloid assay a similar partial rescue was observed, with the central neuroectoderm domain displaying an improved level of compaction, as shown by a smaller area of PAX6+ cells and N-Cad^+^ lumen, when compared to the parental HD cell line (Supplementary Figure S4F and G).

The partial improvement of the HD-signature phenotypes observed in these experiments provides evidence in support of the hypothesis of a dominant negative effect associated with HD mutation.

In the experimental context described above, the inability to fully rescue these phenotypes could be the result of a heterogeneous expression level of the exogenous wtHTT transgene or that the amount of extra wtHTT required to overcome the dominant effect of mtHTT might be required to reach a minimum level to be effective. To address this possibility, we took advantage of the HaloTag included in our transgene constructs (Figure 4A) to enrich for cells homogenously expressing low or high levels of wtHTT-HaloTag by Fluorescence Activated Cell Sorting (Figure 4B). Thus, HD cells complemented with wtHTT-HaloTag were sorted for low levels of HTT (hereafter named as Low), equivalent to endogenous HTT expression level (determined by using a WT knock-in HTT-HaloTag reporter cell line as reference). A population of HD hESC complemented with wtHTT-HaloTag was also sorted for high level of expression of the HTT transgene and hereafter named as High (Figure 4B). HaloTag labelling confirmed that Low and High homogeneously express the transgene at two different levels (Figure 4C), also shown by Immunoblot analysis (Figure 4D). Additionally, linear quantification by MesoScale Discovery Analysis showed that the amount of full length HTT in Low is estimated as expected at ∼2x the endogenous HTT expression level, and High at >10x the level measured in control WT (Figure 4E).

**Figure 4.**
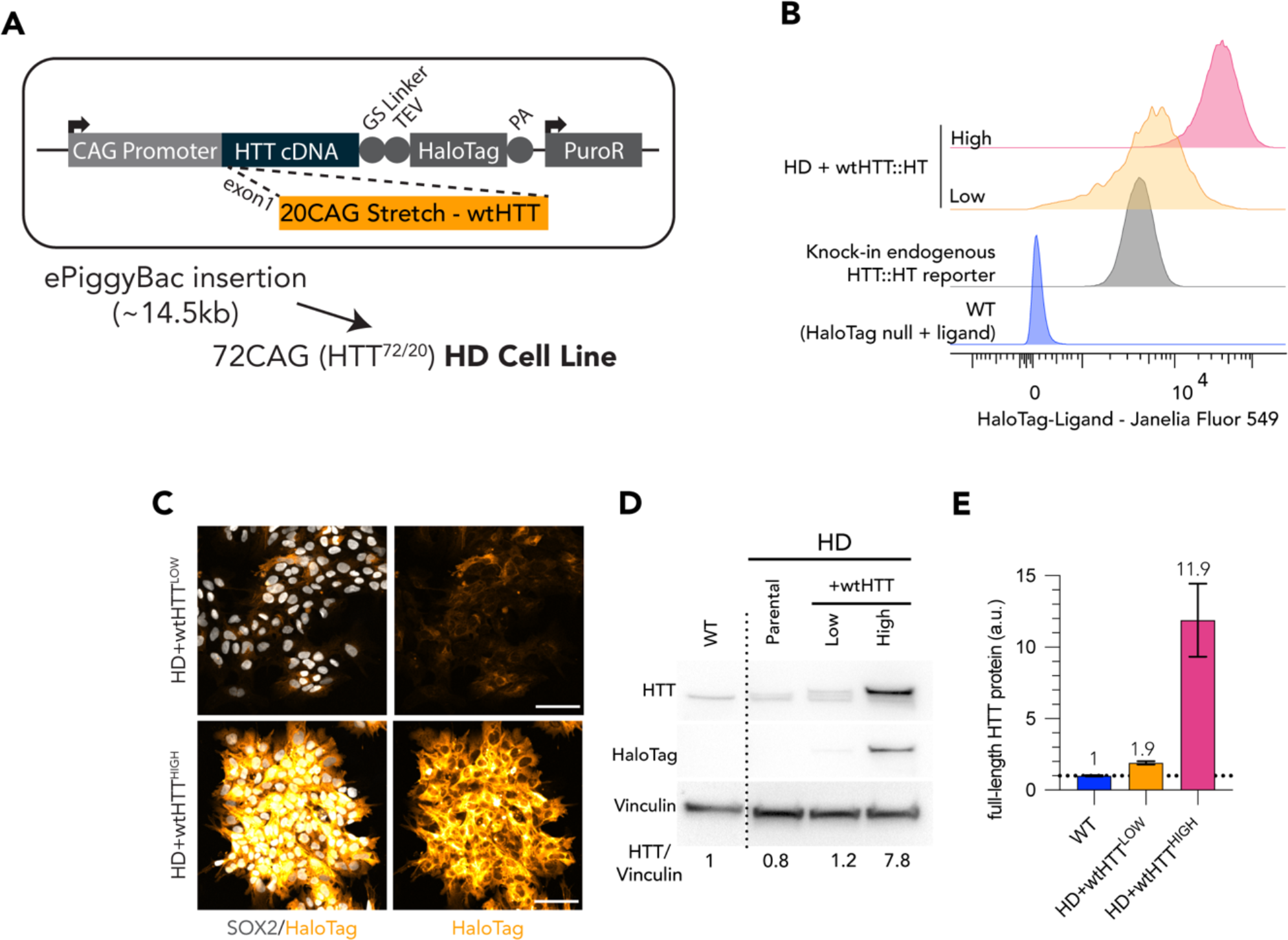
Generation of Low and High wtHTT transgene expression by FACS sorting of live HaloTag fluorescence labelling. **A** Puromycin selectable ePiggyBac transposon carrying full-length wtHTT was inserted into a HD cell line (bi-allelic 20CAG/72CAG background), complementing endogenous expression of wt and mtHTT. **B** Flow cytometry cell counts after fluorescence activated cell sorting of HD+wtHTT cells, into High and Low levels of expression. Low levels were gated using a monoallelic endogenous reporter of HTT with a HaloTag fused into the HTT C-terminal. **C** Chemical tagging of HTT::HaloTag using JaneliaFluor546-Halotag ligand, illustrating homogeneous expression and distinct levels of HTT between Low and High populations. **D** Immunoblot detection of HTT (MAB2166) on HD+wtHTT^LOW^ presents transgene band with exogenous expression of wtHTT::HT at levels comparable to endogenous HTT (larger protein band corresponds to exogenous HTT fused to HaloTag; +36kDa) HD+wtHTT^HIGH^ displays a third band representing the transgene expressed at higher level than endogenous expressed protein. Halotag detection at the HTT molecular weight (∼350kDA) confirms the insertion of the transgene and differential expression levels between low and high fraction. HTT antibody signal quantification (normalized to vinculin, here shown as loading control) is shown below the blot, revealing a ∼0.5X increase and -8.75x increase in comparison to HD background cell line. **F** Linear Quantification of total HTT using Meso-scale discovery assay with 2B7/D7F7 antibody pair (standard curve with polyQ23 full length recombinant human HTT) and 0.9x increase of HTT levels in Low sorted population, and a 11× increase on High expression sorted population. Scale bar: 100µm.

In the HD line complemented with both the Low and High wtHTT expression levels SMAD2/3 nuclear translocation in response to Activin A induction in micropattern cultures resulted in a pronounced WT-like phenotype (p=ns vs. WT cells) (Figure 5A, B). Quantification of the percentage of SMAD2/3 nuclei showed that the number of cells responding to activin A stimulus decreased with over-expression of wtHTT but without reaching the lower level of activated cells observed in WT colonies, regardless of the expression level of the transgene (Figure 5C). Additionally, the degree of rescue was equivalent for Low and High HTT levels, indicating that in the context of this assay complementing an HD cell line with wtHTT only partially rescues the HD phenotype independently of expression level.

**Figure 5.**
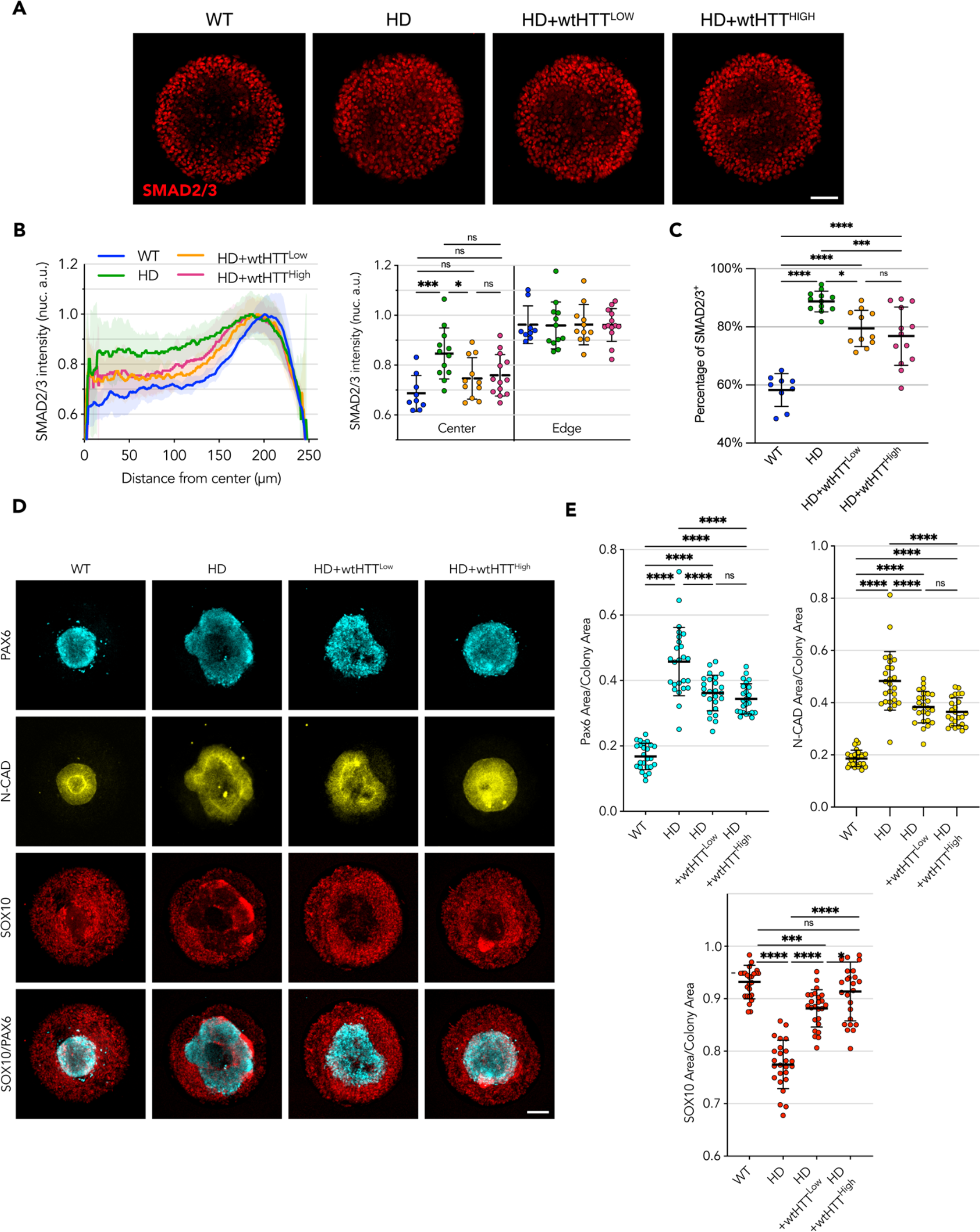
Homogeneous levels of wtHTT transgene expression partially rescue the phenotype in HD cells, independent of expression level. **A** Upon activin A stimulation of 500µm micropatterns, edge restriction of SMAD2/3 nuclear translocation is observed in HD cells complemented with both Low and High levels of wtHTT. **B** Radial profile analysis confirms this behavior, however, not to the levels observed in WT colonies. **D** Binary quantification of positive SMAD2/3 nuclei confirms a trend to amelioration of the phenotypes towards WT-like with increasing wtHTT dose. **E, F** wtHTT resulted in rescued compaction of PAX6/NCAD neuroectodermal domain in the neuruloid assay and recovered SOX10 area. Groups were compared using one-way ANOVA followed by Dunnett’s *post-hoc* test for correction of multiple comparisons (* p<0.05, ** p<0.01, *** p<0.001, ****p<0.0001). All values are presented as mean ± SD. Scale bar: 100µm.

In the neuruloid assay, complementation of the HD cell line with low levels of wtHTT was enough to elicit a rescue towards WT-like phenotype, with improved compact neuroectodermal (PAX6^+^) and N-CAD domains, as well as a larger Neural Crest SOX10^+^ domain than the Parental HD line (Figure 5D, E). On the other hand, complementation with wtHTT expression at high levels resulted in a more pronounced rescue, both in extent of the compaction of the central neuroectoderm domain as well as its morphology, with colonies reaching full closure of the central area and rescue of the area occupied by neural crest cells.

Taken together, these results confirm the hypothesis that, at least partially, the HD signatures observed in a spatially confined pluripotent monolayer in response to activin A stimulation and in the neuruloid self-organization of ectoderm domains are the result of mtHTT that acts in a dominant negative fashion. We show that the CAG-Expanded HTT impairs the normal function of wtHTT, and that mtHTT impairment can be partially rescued with increased expression of wtHTT, although never to a full extent even with levels as high as 10 times greater than the endogenous level.

## Discussion

It has been three decades since HD was first described as a dominant monogenic disease linked to a polymorphic CAG trinucleotide tract in the HTT gene. While research has identified various abnormalities in normal cell and molecular functions as a result of HD mutation (Saudou and Humbert 2016), because the clear function of HTT is still poorly understood, the pathological mechanism resulting from the CAG expansion is still a matter of debate. Due to the presence of numerous confounding factors and the disease’s human-specific nature, traditional approaches like human genetics and phenotypic characterization using animal models have been inadequate in fully resolving this issue. To study HD in humans, our lab generated a genetic allelic series of isogenic human ECS with increasing CAG lengths or HTT deletion (Ruzo et al. 2018) and applied them on an ensemble of micropattern-based assays that model different aspects of human development. Cellular and molecular changes due to HD mutation resulted in highly reproducible and quantifiable functional outputs, with phenotypic severity correlating with CAG length (Galgoczi et al. 2021; Haremaki et al. 2019), as observed with the age of HD onset in human patients.

Although the prevalent hypothesis suggests gain of toxic mechanisms as the cause of HD onset, a growing body of evidence suggests that loss of function might also be a contributing factor (Paine 2015). HTT is ubiquitously expressed from fertilization onwards, with its expression becoming more pronounced in the nervous system (Saudou and Humbert 2016). Lack of HTT expression results in arrested embryonic development at early stage (M. P. Duyao et al. 1995) and conditional deletion of HTT in adult mice forebrain and neurons cultured *in vitro* results in neuronal degeneration, motor phenotypes, early mortality, altered gene transcription and excitatory circuitry (Braz et al. 2022; Dragatsis, Levine, and Zeitlin 2000; McKinstry et al. 2014; Wennagel et al. 2022; Zuccato et al. 2001). Additionally, we found that human HTT-KO ESC lines exhibited *in vitro* phenotypes comparable to CAG-expanded HTT cell lines with identical genetic backgrounds (Galgoczi et al. 2021; Haremaki et al. 2019; Ruzo et al. 2018), suggesting that the HD phenotypes might result of progressive loss-of-function with increasing CAG repeat tract length. Understanding the consequences of HD mutation on the physiological function of HTT is of primary importance for the design of effective clinical approaches for the treatment of the disease and it lacked formal addressing.

In this study, we used a collection of human isogenic cell lines that vary in their HTT genetic makeup (wt/wt, wt/-, -/-, and mt/wt) to systematically investigate the genetic mechanism behind HD phenotypes in previously described *in vitro* micropattern-based assays of activin signaling transduction and self-organized neuronal organoids (Galgoczi et al. 2021; Haremaki, Deglincerti, and Brivanlou 2015). Here, haploinsufficiency (wt/-) did not fully recapitulate the HD-signature phenotypes previously described for HD (wt/mt) and HTT-KO hECSs in these assays (Figure 1), indicating that the molecular insult due to the presence of mutant HTT in our model cannot be explained by a simple loss-of-function mechanism. Similar outcomes in both animal models and human patients were reported previously in other studies indicating that low amount of HTT expression is well tolerated and compatible with normal life. In fact, heterozygote mice with one null HTT allele and one hypomorphic wtHTT allele are embryonically viable and do not present any HD-associated phenotype (Murthy et al. 2019). Furthermore, lowering wild-type huntingtin by ∼45% in the adult non-human primate striatum are phenotypically unremarkable for at least 6 months (Kaemmerer and Grondin 2019). In the human population, HTT levels can vary at least 2-fold or more between different non-HD individuals (Fodale et al. 2022) and although cases of heterozygous individuals with only one functional copy of the HTT gene are rare, they can be found displaying no clinical HD presentation (Ambrose et al. 1994; Jung et al. 2021). Together, these findings are consistent with our results in supporting the idea that although lack of HTT expression results in HD-like outcome, intrinsic mtHTT loss of activity would not be sufficient to elicit the HD-associated phenotypes.

In order to directly determine the effect of HD mutation on HTT function, we overexpressed full length mtHTT in HTT-KO hESCs showing that while it failed to rescue Activin A signaling phenotype in micropatterned cultures, it was able to partially rescue the neuruloid phenotype confirming that indeed HD mutation inactivates (at least partially) the gene function (Figure 2). In agreement with this, it has been reported that HD mutation results in loss of HTT function in neurons (Gauthier et al. 2004). Moreover, other studies have shown that mouse embryonic lethality due to lack of HTT can be rescued with the expression of mtHTT (Leavitt et al. 2001; White et al. 1997), suggesting that this protein retains at least partial function that is sufficient to overcome developmental arrest.

On the other hand, mtHTT transgenic expression in fully competent WT hESCs efficiently induces HD phenotypes indicating that the HD phenotypes observed here are not due to simple inactivation of mtHTT (Figure 3). The HD phenotypes we observe could then be explained as trans-inactivation of wtHTT by the co-expressed mutant HTT, resulting in a HTT-KO-like phenotype due to a dominant negative mechanism, which may have an effect proportional to the CAG repeat lengths (Galgoczi et al. 2021; Haremaki et al. 2019; Ruzo et al. 2018). Consistently, a report on a cis-regulatory variant in the HTT promoter that reduces its expression showed that decreased mtHTT expression delays age of onset while reduction in wtHTT allele expression resulted in earlier onset (REGISTRY Investigators of the European Huntington’s Disease Network et al. 2015).

Considering this putative Dominant negative mechanism, we attempted to rescue the HD phenotype in HD cells by increasing the levels of wtHTT. This resulted in a phenotypic amelioration in both micropattern assays, showing that we are indeed observing a deleterious effect of the mtHTT on wtHTT function that can be overcome by increased wtHTT expression levels. However, by taking advantage of the two highly quantitative micropattern-based assays used, we observed that the phenotypic rescue of HD samples achieved by complementation with additional copies of wtHTT is only partial, even when wtHTT was overexpressed up to 10 times over endogenous levels (Figure 4,5).

Opposing arguments to loss of function mediated by a dominant negative mechanism are that human patients with minimal or no HTT expression (e.g. LOMARS patients) present an extreme juvenile phenotype that differs from the progressive neurodegeneration observed in adult HD patients (Jung et al. 2021; F. Lopes et al. 2016). However, since severe pathology occurs in these patients at early age, it becomes very challenging to discern other brain alterations from complications ahead of normal HD progression. Additionally, even without considering the role of mutant HTT in HD, wtHTT loss is deleterious and therefore total HTT silencing therapies could result in worsened HD phenotype. In that regard, the Phase III GENERATION HD1 (NCT03761849) trial testing an antisense oligonucleotide targeting HTT mRNA, was halted after preliminary analysis revealed that treatment resulted in unfavorable efficacy with worsening HD presentation compared to placebo groups (Schobel 2021). While neuroinflammation or other adverse response to the therapeutic agent could also explain this unfortunate outcome, one must consider the possibility of a dominant negative effect and therefore interpret the observed worsening of HD clinical manifestation as the consequences of lowered total HTT levels in HD patients, resulting in accelerated HD progression.

Overall, although we cannot exclude a component of pure gain of toxicity by mtHTT might also contribute to HD pathogenesis, we show here by corollary that the hindering of HTT physiological function by mtHTT (both in cis and trans) is at play in our system, and that the HD-associated phenotypes can be improved by complementation with additional copies of wtHTT, strongly supporting a classical dominant negative mechanism. A clear understanding of the molecular mechanisms behind the phenomenon that we described here is still lacking. However, because we could not fully rescue the phenotype with wtHTT overexpression, we speculate that the dominant negative mechanism observed could be the result of wtHTT – mtHTT crosstalk (either direct or indirect). Future research will have to address how wtHTT biology is changed in the presence of its mutant counterpart, including but not limited to its interaction partners and its subcellular localization, which will provide the HD field with additional therapeutic targets to cure, ameliorate or halt progression of HD pathology. Until such advances are realized, increasing wtHTT levels could provide some degree of protection to HD patients. Alternatively, our results suggest that the most promising therapeutic approach at this stage should aim at allelic-specific silencing of the mtHTT gene or targeted ablation of poly-glutamine expanded HTT protein, despite the technical challenges to implement this universally.

## Methods

### Human ES Cell Culture

Cell lines used in this study were derived from the RUES2 cell line, part of the NIH Human Embryonic Stem Cell Registry (RUES2-NIHhESC-09-0013). hESCs were maintained in mouse embryonic fibroblast conditioned HUES Media in feeder free conditions (conditioned media – CM), supplemented with bFGF 20 ng/mL, and replaced daily. hESCs were passaged at 70-80% confluency every 3-4 days into Geltrex coated dishes (Life Technologies) using gentle cell dissociation reagent (STEMCELL-Technologies). Cells were tested for *Mycoplasma* spp. at 2-monthly intervals.

### Stable integration of full length HTT-HaloTag piggyBac vectors in hESCs

Human ESCs were nucleofected using an Amaxa nucleofector II in Nucleofector Solution L (all Lonza) using B-016 preset. A total of 106 hESCs were nucleofected with 1µg of piggyback HTT-HaloTag plasmid (wtHTT or mtHTT according to each experiment) and 1µg of the piggyBac transposase (Lacoste, Berenshteyn, and Brivanlou 2009). Cells were plated on Geltrex coated plates in CM supplemented with bFGF 20 ng/mL and 10 µM ROCK-inhibitor (Y-27632) for 24h. Cells were then selected with puromycin 1ug/ml for up to 10 days and expression validated by and in situ labelling of HaloTag, flow cytometry, and immunoblot to confirm HTT-Halotag fused transgene.

### Immunoblotting

Protein was extracted in ice using ice cold RIPA Lysis and Extraction Buffer (Thermo Scientific 89900) supplemented with 1× Halt™ Protease and Phosphatase Inhibitor Cocktail (Thermo Scientific 78441). Cell lysates were briefly sonicated using a Branson Sonifier 450 (VWR; output 4, duty-cycle 10%, allowing 5 bursts) and incubated on ice for 15 min. Lysates were cleared by centrifuging at 12.000 rpm for 5 min at 4°C and protein concentration determined using the Pierce™ BCA Protein Assay (Thermo Scientific 23225). Samples were prepared for SDS-Page in 1× NuPAGE™ LDS Sample Buffer containing 50mM DTT, boiled at 95°C for 5 min and loaded into a NuPAGE 3-8% Tris-Acetate Gel (Thermo Scientific) along with HiMark™ Pre-stained Protein Standard (Thermo Scientific LC5699). Electrophoresis was run on ice and under stirring for 1h30 at 150V. Protein bands were transferred to a Trans-Blot Turbo 0.2um PVDF membrane (Bio Rad Cat.#: 1704156) using a Trans-Blot Turbo Transfer System (Bio-Rad) and membranes blocked in Tris-buffered saline containing 0.1%Tween-20 (TBS-T) and 5% milk for 30min. Primary antibodies were incubated ON at 4°C and HRP secondary antibodies for 30 min at RT. All incubations were carried under constant agitation and washed thoroughly between steps. HRP was detected using either Clarity or Clarity Max ECL Western Blotting Substrates (Biorad) and chemiluminescence detected in a Chemidoc MP system (Biorad).

### Activin A 2D Micropattern Assay

Spatially confined hESC response to activin A was performed as previously described(Galgoczi et al. 2021). Briefly, custom micropatterned glass coverslips presenting 500 µm groWTh areas (CYTOOCHIPS Arena 500A) were covered with 800µL of 10 μg/ml recombinant human laminin 521 (BioLamina) diluted in PBS^+/+^(Gibco) for 3h at 37°C. Coated coverslips were transferred to a 35mm dishes containing 5mL of PBS^+/+^ and serially washed 5 times, avoiding full removal to avoid drying, followed by one final complete wash with PBS^+/+^. Cells lines were precultured in parallel to ensure similar growth upon the start of the assay. hESCs were dissociated in single cells using accutase (Stem Cell Technologies) and 0.8×10^6^cells seeded per micropattern in 3mL of CM with 20 ng/ml bFGF and 100 μg/ml Normocin (InvivoGen).

ROCK inhibitor, Y27632 (10 μM; Abcam ab120129). Micropatterns were left unperturbed for 10 min to ensure homogenous distribution across the patterns and then transferred to 37°C. ROCK inhibitor was removed 3h after seeding and upon 18h incubation, the monolayers were induced with 100ng/ml of Activin A (R&D Systems). After 1h incubation, samples were washed, fixed, and processed for Immunofluorescence.

### Neuruloid Micropattern Assay

Neuruloid culture (Haremaki et al. 2019) was performed in coated 500µm Micropatterns as described above. Briefly, 0.5×10^6^cell were seeded per micropattern and incubated for 3 h at 37°C in HUESM medium supplemented with 20 ng/mL bFGF, 10 μM Rock inhibitor Y27632 and 100 μg/ml Normocin. Cultures were kept in Normocin for the whole experiment. The seeded micropatterns were then washed once with PBS^+/+^ and day 0 initiated in HUESM with 10 μM SB431542, 0.2 μM LDN 193189. On day 3, media was replaced with HUESM containing 10 μM SB431542 and 3 ng/mL BMP4, and media replenished at day 5. Micropattern cultures were then fixed at Day 7 and processed for Immunofluorescence analysis.

### Immunofluorescence

Cells were fixed with 4% paraformaldehyde (Electron Microscopy Sciences 15713) for 30 min at RT, washed trice with DPBS and permeabilized and blocked DPBS containing 0.5% Triton-X 100 (Sigma 93443) and 3% Normal Donkey Serum for 30 minutes. Primary antibody staining was performed in DPBS with 0.1%TX-100 (PBST) for 1h30 at room temperature, washed trice with PBST for 5 min each, and secondary antibody staining was performed for 1h in PBST at RT. After 2 washes with PBST and 1 PBS wash, samples were counterstained with 0.1 µg/mL DAPI (Thermo Fisher Scientific D1306) for 10 min and washed. Micropattern coverslips were mounted on slides using ProLong Gold antifade mounting medium (Molecular Probes P36934). hESCs grown in IBIDI slides were mounted in IBIDI mounting medium and stored at 4 °C before imaging. Antibodies used can be found in Table S1.

### Fluorescence Activated Cell Sorting (FACS)

Cells grown under confluency were incubated in CM+bFGF media containing 1μM cell permeable Janelia Fluor® 549 HaloTag ligand complex (Promega, GA1110) for 1 hour at 37°C. Cells were washed 3 times with PBS^+/+^ and incubated with fresh media to allow unbound ligand to be fully removed for 3 hours. Monolayers were washed again with PBS^-/-^, dissociated with accutase and resuspended in PBS -/-, containing HEPES 10 mM, EDTA 5 mM, BSA 0.5%. Duplets were excluded based on FSC-W and SSC-W gating and DAPI was used at XX µg/mL for cell death exclusion. Cells were sorted in a FACS Aria Flow Cytometer (BD) using a 100µm/20psi nozzle. An endogenous C-terminal tagged CRISPR HTT reporter with HaloTag (unpublished) was used to define the gating for Low levels of HTT, and high levels were defined as all cells above that threshold. The parental untagged cell line was treated with HaloTag ligand in parallel and used as negative control. Cells were allowed to recover after sorting in Normocin and the obtained populations analyzed by flow cytometry, Western Blot and HTT MSD quantification.

### Electrochemiluminescence Meso Scale Discovery for HTT expression quantification

To quantify total HTT expression levels on the generated cell lines, about 3×106 cells were pelleted and flash frozen in triplicates and sent for contracted analysis at Evotech with MSD assay CHDI_HTT_144 (antibody pair 2B7/ D7F7) following SOP. Briefly, samples were lysed in ice cold MSD Tris Lysis Buffer and incubated 30 min at 4°C by rotating. After lysis, the samples were centrifuged (15700 rcf, 4°C; 6 min) and protein concentration determined with BCA assay and adjusted to 0.2 mg/mL, further diluted to 0.1mg/mL in blocking buffer. Final buffer conditions for HTT quantitation assays were 20 mM Tris (pH 7.5), 150 mM NaCl, 1 mM EDTA, 1 mM EGTA, 1% Triton-X, 1× phosphatase inhibitor I and II (Sigma), complete mini EDTA free protease inhibitor cocktail tablet (Roche), 0.5 mM PMSF, 10 mM NaF. Purified recombinant full length 1-3144 protein of human Huntingtin protein with 23 polyglutamine(CHDI-90001858) was used as standard, ranging between 0.001 and 10 fmol/µL.

### HaloTag in vitro ligand Staining

For imaging, hESCs were grown in glass surfaces previously coated with 10μg/ml recombinant human laminin 521 (BioLamina) diluted in PBS^+/+^(Gibco) for 3h at 37°C. Freshly thawed Janelia Fluor® 549 HaloTag® Ligand (Promega, GA1110) was diluted to 200nM in warmed CM and cells incubated for 30 min. After washing thrice with PBS^+/+^, cells were incubated for 30 min to allow unbound ligand to wash. Cells were then fixed in 4% PFA, nuclei counterstained with DAPI and imaged by confocal microscopy.

### Imaging

Confocal microscope images of hESCs and neuruloid IF, were acquired with a 10x or 20×/0.8 NA dry objective lens (Zeiss), using 405, 488, 561 and 633 nm laser lines, and a combination of PMT and GaAsP detectors (LSM 780, Zeiss). Large tilled imaged were obtained to sample the neuruloid assay. Activin A assay was imaged acquiring large tiled areas with an inverted wide-field epifluorescence microscope using 10×/0.45 NA dry objective lens (Zeiss Axio Observer Z1).

### Image analysis

Fiji (2.0.0/1.53t) was used to stitch tilled images and to generate maximum projections of the generated confocal z-stacks. SMAD2/3 radial intensity profiling was performed as previously described with slight changes (Galgoczi et al. 2021). Briefly, using a custom Phyton script, a foreground mask was created to detect colonies by thresholding the DAPI channel and calculating alpha shapes with respect to predicted colony size, and colonies extracted into individual files. Ilastik (1.4.0b27, Berg et al. 2019) was then used to segment individual nuclei based on the DAPI channel. Identified nuclei were then filtered according to expected size to exclude debris and mitotic cells, and the center of the nuclear mask was determined for further watershed segmentation (scikit-image). The generated nuclear mask was then applied to the SAMD2/3 channel and median nuclear intensities were extracted, and SMAD2/3 radial profile was plotted according to each nucleus distance to the center of the colony. Profiles were normalized to 1 using the maximum value obtained in the profile.

To assess the total SMAD2/3 number, individual nuclei were segmented using Stardist (Schmidt et al., 2020; https://arxiv.org/abs/1806.03535), and the corresponding nuclear masks were used to calculate the mean intensity for each individual nucleus (scikit-image). Images were manually thresholded for SMAD2/3 to classify nuclear translocation as binary. The percentage of SMAD2/3 positive nuclei was then calculated on the total nuclei number.

Individual neuruloid images were obtained as explained above. Ilastik was used to segment the areas occupied by each marker and DAPI was used to determine the total neuruloid area. The pixel area for each marker was then determined and normalized as a fraction of the total DAPI area. Python libraries numpy, pandas, matplotlib and seaborn were used to organize the data and for plotting.

### Statistics

All data presented in this study were obtained from at least two independent experiments. Statistically significant differences between conditions were determined using one-way analysis of variance (ANOVA) followed by Dunnett’s post-hoc test for correction of multiple comparisons. Statistical analyses were performed in Prism 9 (GraphPad). Results were considered not significant (n.s.) when the observed differences when p>0.05, and significance levels were denoted as follows: *p<0.05, **p<0.01, and ***p<0.001.

## Acknowledgments

We extend our sincere appreciation to the members of the Brivanlou lab for their valuable contributions and insightful discussions during this research. We are grateful for the support provided by the Rockefeller University Bio-Imaging Resource Center, particularly Alison North and Christina Pyrgaki, who offered exceptional assistance in microscopy. Special thanks are also due to the Flow Cytometry Resource Center and its team, including Svetlana Mazel, Stanka Semova, Samer Shalaby, and Songyan Han, for their technical expertise and guidance with the flow cytometry experiments. We acknowledge the early work on the full-length cDNA HTT piggyback plasmid by former lab members Albert Ruzo and Jeffrey Naftaly. We are also grateful to Jakob Metzger for sharing and adjusting single cell quantification image analysis tools used for Activin A radial quantification experiments. Additionally, we would like to express our appreciation to Tom Vogt, Dan Felsenfeld and David Howland from the CHDI Foundation for their valuable support and advice throughout this project.

## Author Contributions

TL: study design, data acquisition and analysis, writing original draft. SL, EC: data acquisition. RdS: data analysis, review, and editing. FMP: conceptualization, study design, supervision, writing original draft, review, and editing. AHB: conceptualization, funding, supervision, review, and editing. All authors contributed to the article and approved the submitted version.

