## Supplementary Figures for "HD mutation results in a dominant negative effect on HTT function"

### SUPPLEMENTARY FILES

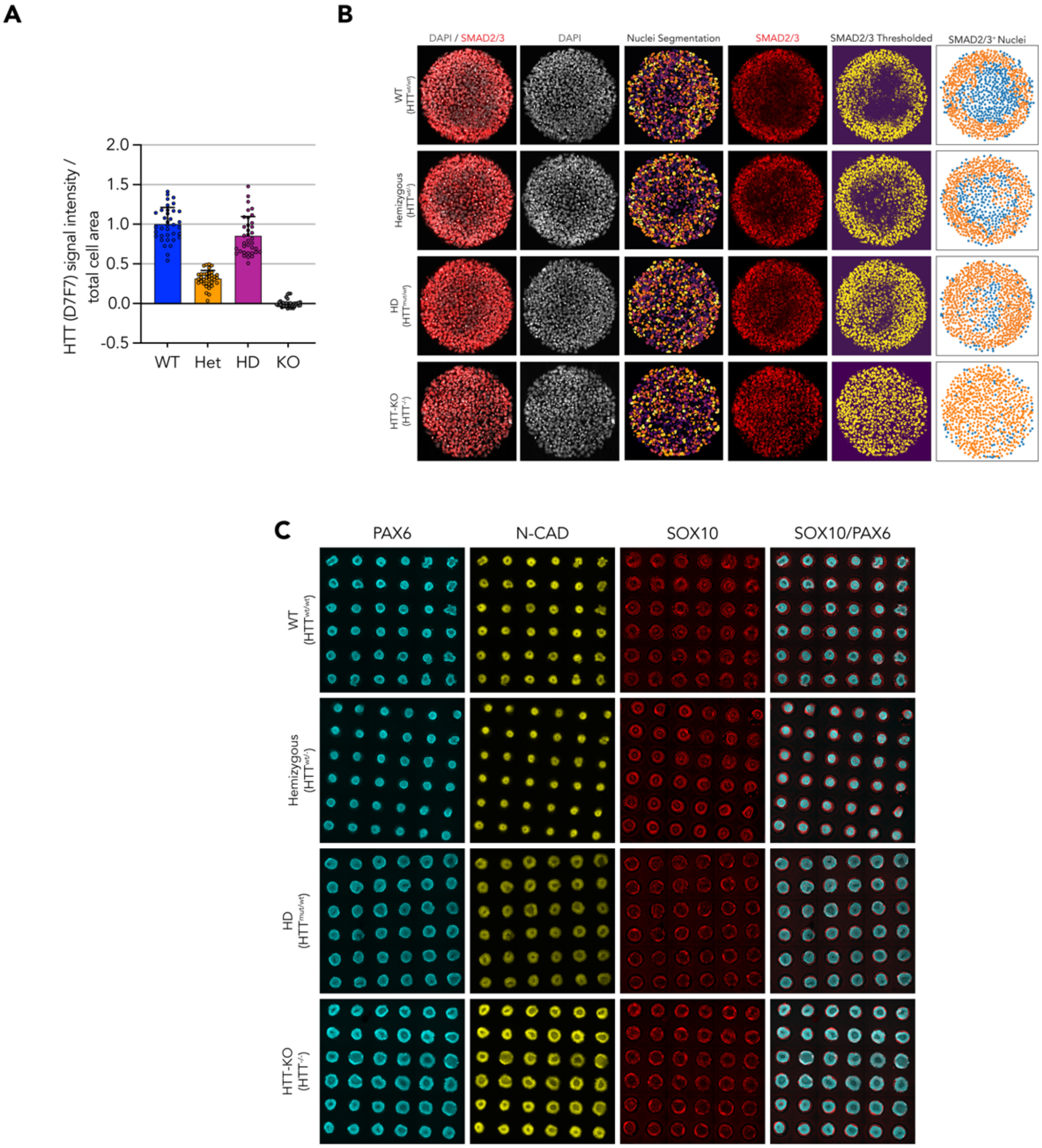


**Figure S1 A** Quantification of immunofluorescent signal of HTT (D7F7 antibody) reveals halving of HTT amount in the Heterozygous cell line, compared to WT and HD cell Line; signal density was normalized to area occupied by ESC colonies (n=36 fields). **B** Pipeline for quantification of Fraction of SMAD2/3$^+$ nuclei; Segmented individual nuclei were matched with their integrated SMAD2/3 density and classified as positive (orange) and negative (blue) after manual thresholding. **C** Overview of the micropatterned cultures used in the neuruloid assay show the range of variability between individual colonies observed in this assay.

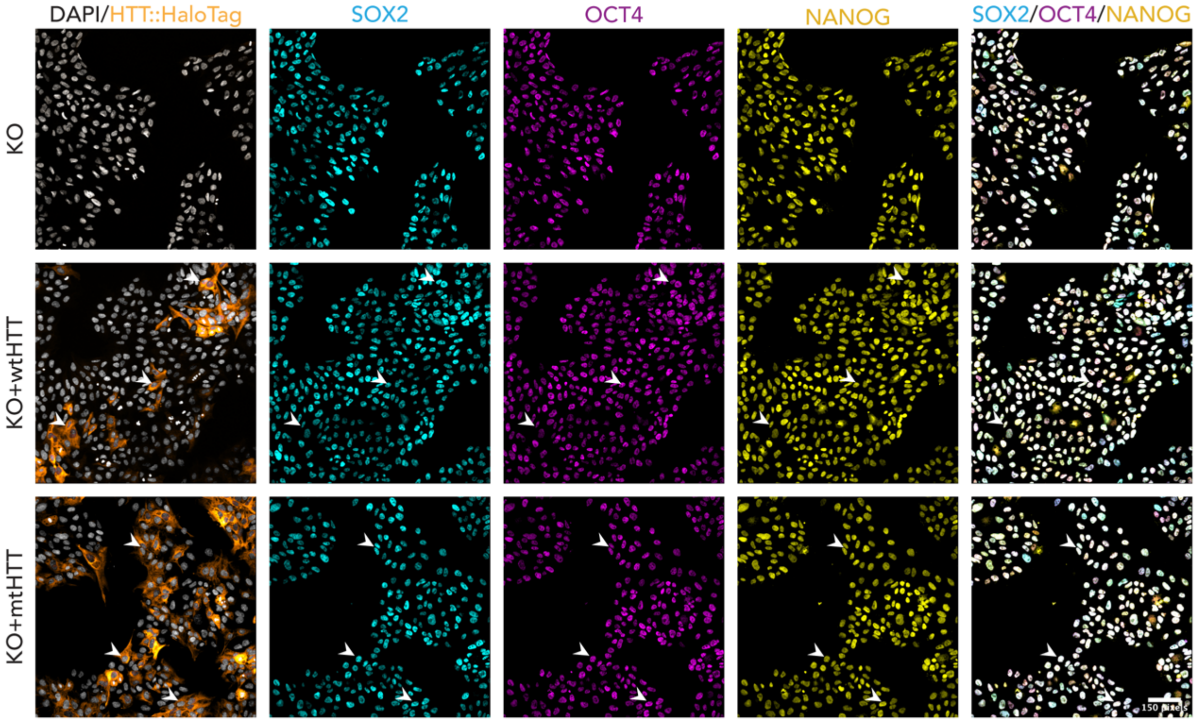


**Figure S2 -** HTT-KO cell line carrying wt or mtHTT-HaloTag transgene retain co-expression of pluripotency markers SOX2, OCT4 and NANOG. Scale bar: 100$\mu$m.

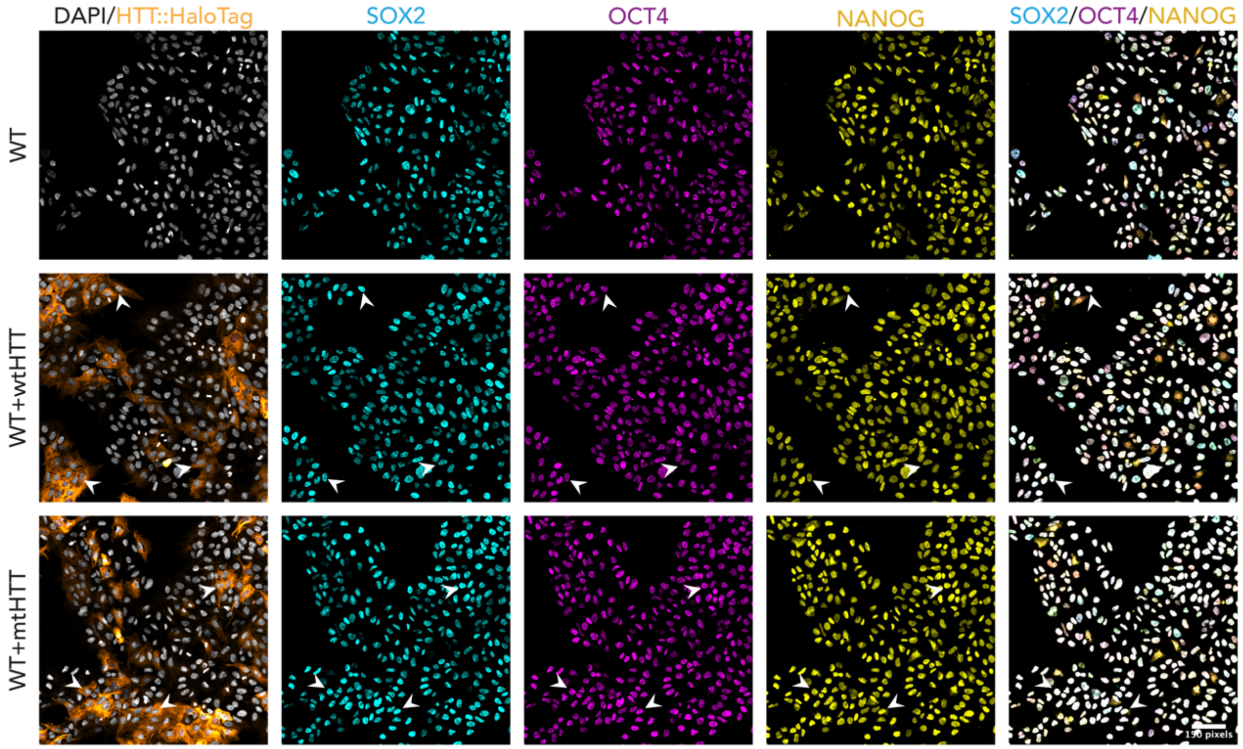


**Figure S3 -** WT cell lines carrying wt or mtHTT-HaloTag transgene complementing endogenous HTT levels retain co-expression of pluripotency markers SOX2, OCT4 and NANOG. Scale bar: 100$\mu$m.

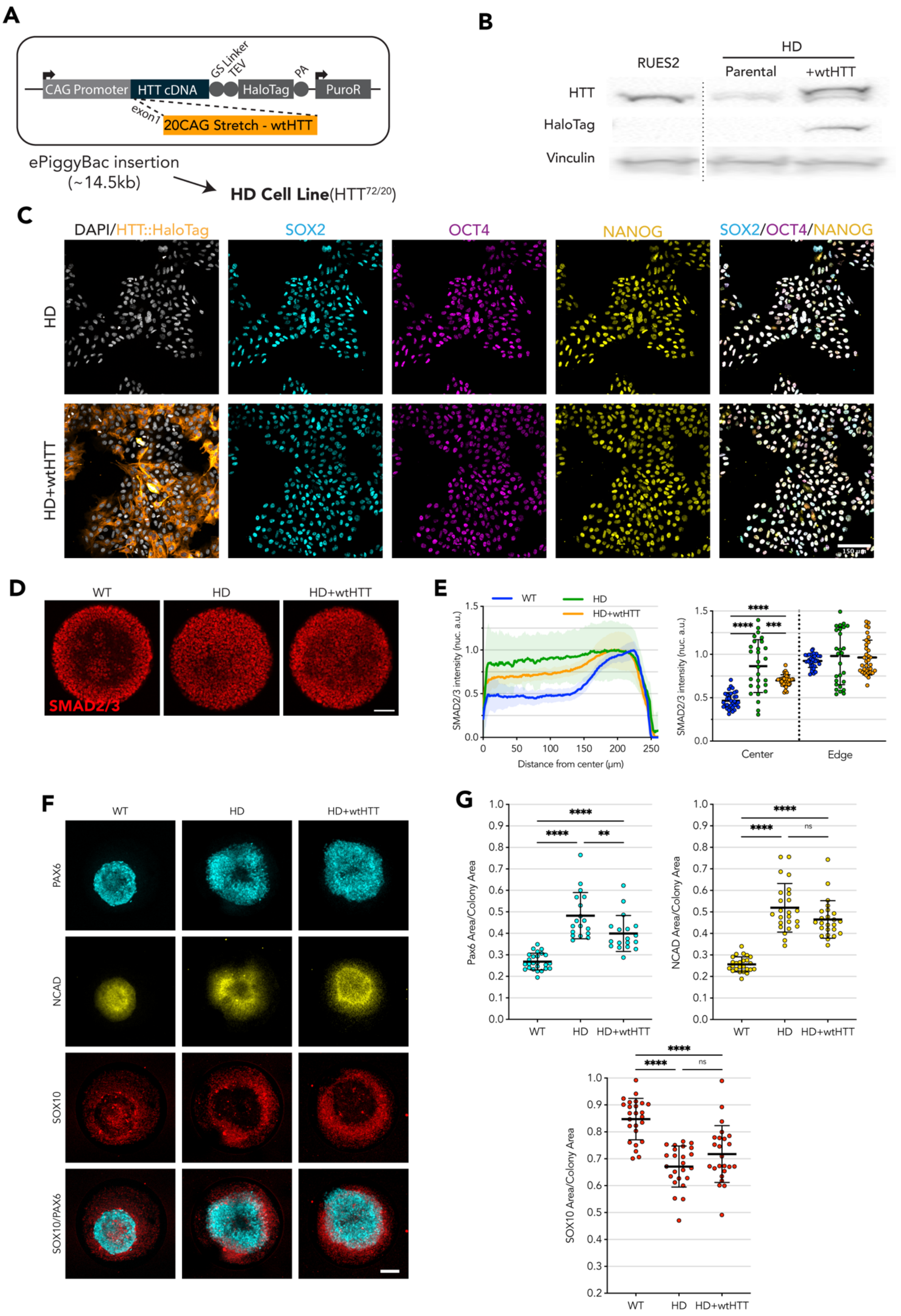

A
CAG Promoter
HTT cDNA
GS Linker
TEV
HaloTag
PA
PuroR
exon1
20CAG Stretch - wtHTT
ePiggyBac insertion
(~14.5kb)
HD Cell Line(HTT72/20)
B
RUES2
HD
Parental
+wtHTT
HTT
HaloTag
Vinculin
C
DAPI/HTT::HaloTag
SOX2
OCT4
NANOG
SOX2/OCT4/NANOG
HD
HD+wtHTT
D
WT
HD
HD+wtHTT
SMAD2/3
E
WT
HD
HD+wtHTT
SMAD2/3 intensity (nuc. a.u.)
Distance from center (μm)
Center
Edge
F
WT
HD
HD+wtHTT
PAX6
NCAD
SOX10
SOX10/PAX6
G
Pax6 Area/Colony Area
NCAD Area/Colony Area
SOX10 Area/Colony Area
WT
HD
HD+wtHTT
ns

**Figure S4 – wtHTT transgene expression partially rescue the phenotype in HD cells A** Puromycin selectable ePiggyBac transposon carrying full-length wtHTT was inserted into a HD cell line (bi-allelic 20CAG/72CAG background), complementing endogenous expression of wt and mtHTT. **B** Immunoblot against HTT and HaloTag reveals the stably integrated wtHTT-HaloTag construct in the Parental HD line. **C** HD ($HTT^{20/72}$) cell lines carrying wtHTT-HaloTag transgene complementing endogenous HTT levels retain co-expression of pluripotency markers SOX2, OCT4 and NANOG. **D** Immunofluorescence imaging shows SMAD2/3 nuclear translocation upon Activin A stimulation, limited to the edges on WT and Het colonies with wider to full induction on the colony center HD and to a lesser extent in HD+wtHTT colonies. **E** Mean radial intensity profile of nuclear SMAD2/3 shows insensitivity to Activin A at the colony center in HD+wtHTT is rescued, albeit partially (scatter plot displays the mean SMAD2/3 nuclear intensity for each colony at center:25-10$\mu$m and edge:175$\mu$m-225$\mu$m; WT n=34, HD n=28, HD+wtHTT n=36). **F** Representative immunofluorescence images of Neuruloid induction assay performed on 500$\mu$m circular micropatterns. **G** Complementation of HD hESCs with wtHTT transgene resulted in a reduction of PAX6 area, showing partial rescue of the phenotype (WT n=25, HD n=24, HD+wtHTT n=24). Groups were compared using one-way ANOVA followed by Dunnett's *post-hoc* test for correction of multiple comparisons (* $p<0.05$, ** $p<0.01$, *** $p<0.001$, ****$p<0.0001$). All values are presented as mean ± SD. Scale bar: 100$\mu$m.

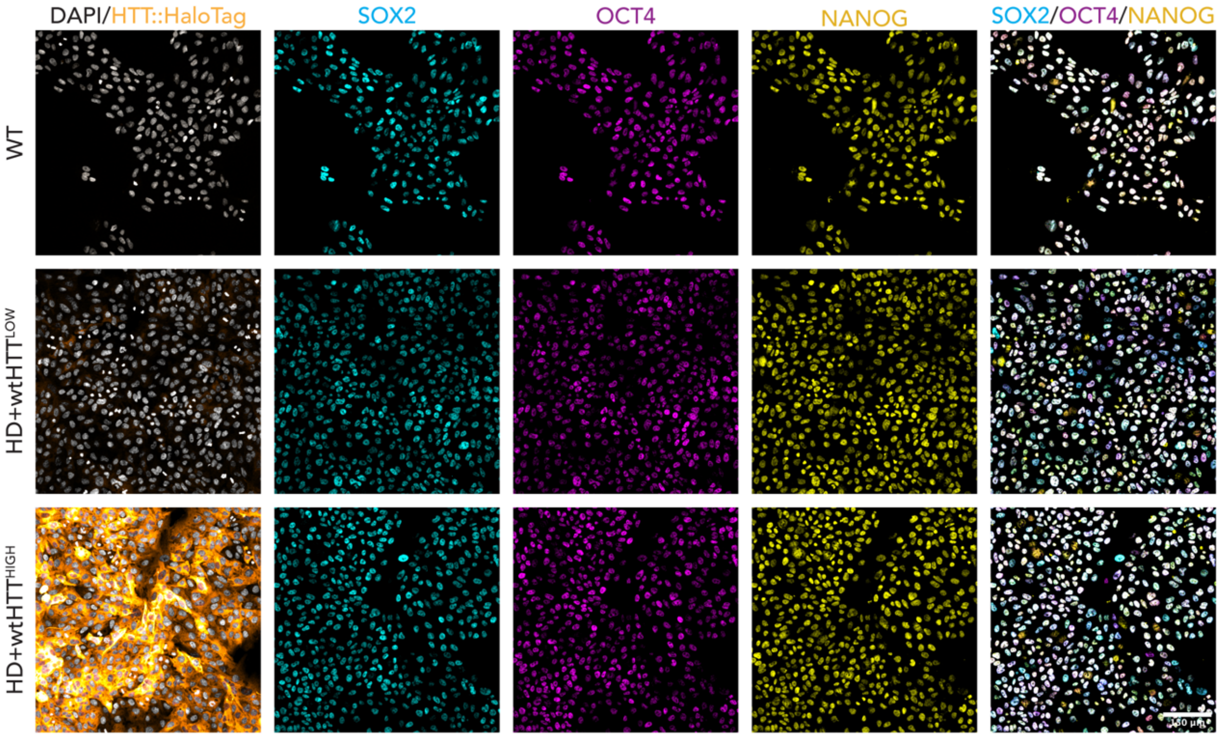


**Figure S5 –** HD ($HTT^{20/72}$) cell lines carrying wtHTT-HaloTag transgene at Low and High levels after FACS, retain co-expression of pluripotency markers SOX2, OCT4 and NANOG. Scale bar: 100$\mu$m.
